# *Plasmodium* blood-stage induces trained immunity in hepatocytes

**DOI:** 10.64898/2026.07.20.739619

**Authors:** Elizabeth KK Glennon, Tinotenda Tongogara, Nhi Ho, Veronica I Primavera, Noah Stegman, Ling Wei, Lola Velush, Cassandra Fieldson, Bogdan Velychko, Ashley Subijanto, Rafael B. Polidoro, Sean Murphy, Raj P Kapur, Nana K Minkah, Nathan W Schmidt, Tuan M Tran, Alexis Kaushansky

## Abstract

*Plasmodium* parasites, the causative agents of malaria, are transmitted at high levels in endemic areas and sequential infections are common. Using a mouse model of infection we discovered liver burden was suppressed in blood-stage experienced, compared to naïve, animals, independently of adaptive responses and inflammation. We observed greater chromatin accessibility of a subset of interferon stimulated genes in blood-stage experienced animals, and a rapid increase in transcription of these genes upon sporozoite challenge. *Ex vivo* stimulation of hepatocytes from blood-stage experienced mice also led to a rapid and elevated response upon treatment with an unrelated antigen, ctDNA. Taken together, these data are consistent with a model in which hepatocytes are reprogramed by blood stage *Plasmodium* infection to exhibit trained memory akin to what has been described in innate immune cells. The consequences of training of hepatocytes could be wide-reaching and might alter responses to diverse pathogens and other stimuli.

## Introduction

Infection dynamics are typically studied in isolation in the laboratory, whereas in the field pathogens occur, recur and co-occur. When multiple infections are evaluated in the laboratory, it is typical to assess the adaptive, antigen-dependent phenomena that control infection and pathogenesis. Yet, in many cases, these laboratory observations do not entirely explain observed outcomes or the epidemiology in the field. Innate immune memory describes an epigenetic-driven, antigen-independent reprograming of innate immune responses to sequential infections (*1*). Growing evidence shows innate memory plays a role in regulating both infection and immunity (*2, 3*).

Malaria infection and pathogenesis exemplify the complex dynamics of sequential infection that can occur in the field. Individuals are regularly infected with *Plasmodium* parasites, experiencing hundreds of infectious mosquito bites a year in some endemic regions (*4, 5*). These high levels of exposure, however, do not result in sterile immunity. Instead, adults in malaria-endemic areas develop partial immunity (control of parasitemia) and tolerance (reduced disease symptoms), leading to asymptomatic cases with prolonged low or even submicroscopic parasitemia (*6–8*). It remains largely unknown how these complex dynamics impact the progression of the malaria life cycle.

Upon transmission via the bite of a mosquito, the *Plasmodium* sporozoite migrates to the liver where it undergoes development within a hepatocyte before moving to the blood stream. Once in the blood, subsequent cycles of parasite replication in red blood cells (RBCs) lead to disease. Recent studies have begun to describe the capacity of *Plasmodium* host-pathogen interactions to influence pathogenesis across parasite stages, independently of adaptive immune responses, both within a single infectious challenge or in the context of superinfections occuring in quick succession (*9–11*). The dynamics of liver-stage (LS) infection for instance, can influence the development of blood-stage (BS) pathogenesis, including cerebral malaria, within the same infection (*10, 12*). Studies to date on sequential infections in the field have indicated a role for innate memory of circulating monocytes in malaria-experienced individuals. However, the nuances of the responses have varied across studies, with some groups describing the development of a tolerized state and others an enhanced immune response to secondary stimulus suggestive of trained immunity (reviewed in (*13*)).

Here, we use a sequential infection mouse model of chronic BS malaria with prolonged submicroscopic parasitemia, followed by LS infection with a congeneric malaria species, to assess the antigen-independent tuning of memory responses that shape the infection trajectory during *Plasmodium* sporozoite challenge in non-naïve animals. We have selected this approach to model what occurs in the field, where a new infection often follows the previous one, which has often not entirely cleared (*14, 15*). We observe a reduction in LS infection in BS-experienced animals that is independent of adaptive T-cell and antibody responses, and evidence that BS infection epigenetically reprograms the liver to a state where IFN-responsive genes are more accessible and more rapidly transcribed in hepatocytes in response to a secondary stimulus. Our data are consistent with the hypothesis that hepatocytes can exhibit innate immune memory which, to our knowledge, has not previously been reported in a mammalian system. It remains an open question to what extent *Plasmodium* training of hepatocytes might alter the trajectory of other infections and under what other conditions trained immunity is elicited in hepatocytes.

## Results/Discussion

To investigate how prior exposure to BS *Plasmodium* influences subsequent LS infection in an antigen-independent manner, we employed a sequential infection model with two rodent-infectious *Plasmodium* species. We infected mice with *P. chabaudi* BS parasites, bypassing LS infection, and monitored their parasitemia until it was submicroscopic, between 20-22 days post-infection (dpi) (Fig. 1A, fig. S1A). A majority of these animals retained infection as monitored via quantitative PCR (qPCR) (fig. S1B). We then infected *P. chabaudi* experienced mice, or naïve controls, with *P. yoelii* sporozoites via retro-orbital injection. Mice were necropsied and livers were isolated after 44 hours, when the parasites had progressed to late LS infection (Fig. 1A). *P. chabaudi* BS experienced mice had significantly lower LS burden 44 hours after *P. yoelii* sporozoite challenge compared to previously naïve mice, as measured by levels of *Py*A18S transcript in the liver (Fig. 1B, fig. S1C). Schizont size did not differ significantly between the two groups, indicating the difference was due to fewer, rather than smaller, parasites (fig. S1D).

**Fig. 1.**
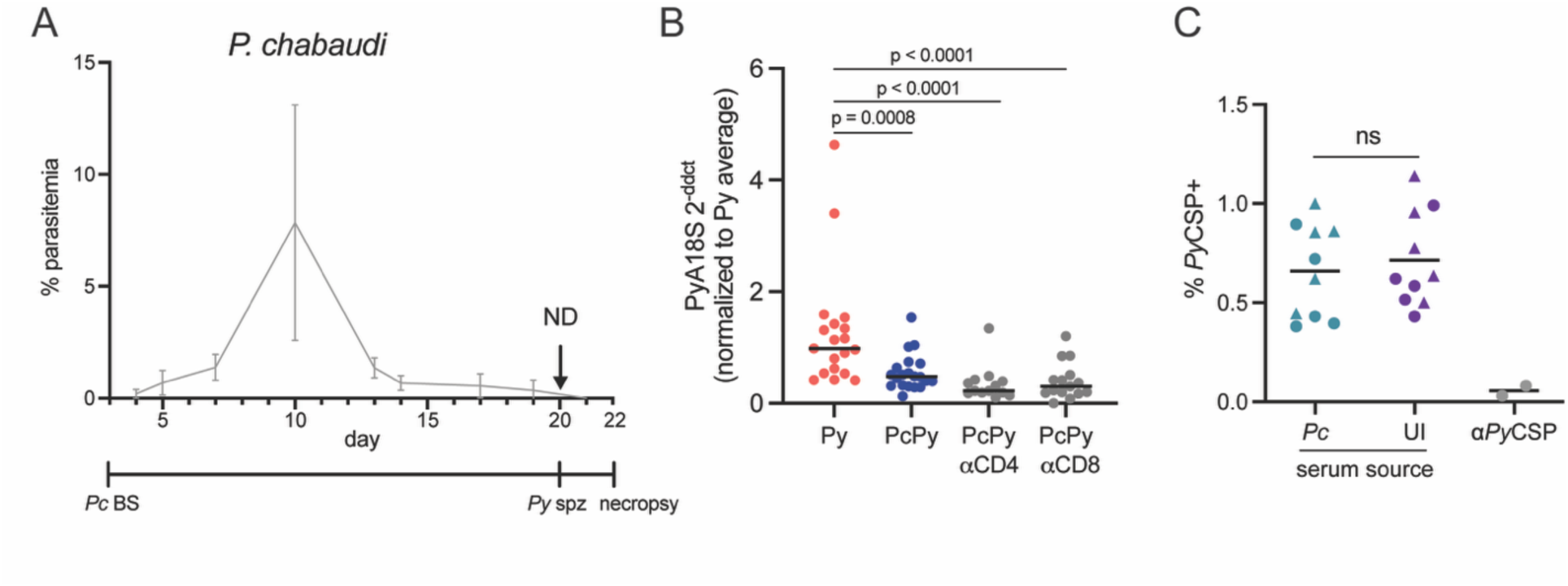
*P. **chabaudi*** **blood-stage infection protects against subsequent *P. yoelii* liver-stage infection independently of CD4^+^ or CD8^+^ T-cells or antibody-mediated effects on invasion. (A)** Schematic of experimental timeline. Mice were injected with *P. chabaudi* blood-stage (*Pc* BS) parasites on day 0 and parasitemia monitored by blood smear. Once BS parasites were not detectable by microscopy (ND), around day 20, mice were infected with *P. yoelii* sporozoites (*Py* spz) and then necropsied at 44hpi. **(B)** PyA18S quantified by qPCR in livers of mice 44h post-sporozoite infection. Mice were either naïve (*Py*) or had previously been infected with *P. chabaudi* BS parasites for 20-26 days (PcPy) and had CD4^+^ or CD8^+^ T cells depleted by antibody (αCD4 or αCD8, respectively) injection before sporozoite injection. Data were normalized to Gapdh and to the average of control (*Py*) mice within each experiment. Each dot represents an individual mouse from one of 4 independent experiments. Data were analyzed by the Mann-Whitney test. **(C)** Percentage PyCSP^+^ hepatocytes measured by flow cytometry 90 minutes after infection by *P. yoelii* sporozoites. Sporozoites were incubated with serum from male or female uninfected (UI) or day 20 *P. chabaudi*-infected (*Pc*) mice before invading Hepa1-6 cells for 90 minutes. Each dot represents cells infected with sporozoites treated with serum from an individual mouse or 0.46mg/mL 2F6 antibody as a positive control.

Antigen-specific CD8^+^ T cells have been described to be potent regulators of LS infection (*16*). To assess the role of the T cell response in this system, we conducted the same sequential infection experiment but depleted CD4^+^ or CD8^+^ T cells by antibody injections administered after *P. chabaudi* parasitemia resolved and prior to *P. yoelii* sporozoite challenge (fig. S1E-G). The suppressive effect of *P. chabaudi* BS infection on LS infection was not reversed by depletion of either T cell population (Fig. 1B). In addition to T-cells, several studies have demonstrated the capacity of sporozoite-specific antibodies to block LS infection (*17*). While unlikely, given previous studies demonstrating antibody-based protection is not conserved between *P. chabaudi* and *P. yoelii* (*18*), the observed suppression of LS burden could be caused at the point of invasion by cross-reactivity of the humoral adaptive response induced by *P. chabaudi* BS with *P. yoelii* sporozoites. To test this possibility, we incubated *P. yoelii* sporozoites with serum collected from *P. chabaudi*-experienced mice 20 dpi and saw no difference in hepatocyte invasion rates *in vitro* compared to parasites treated with serum from uninfected mice (Fig. 1C). Incubation with the positive control anti-PyCSP antibody 2F6 reduced invasion rates as expected (Fig. 1C)(*19, 20*).

The liver plays a role in clearing parasitized RBCs and is a major site of inflammation during BS infection (*21*). We reasoned that residual inflammation or resulting pathology in the liver, or a prolonged systemic cytokine response, could suppress LS burden upon secondary challenge. To assess this possibility, we further characterized the liver microenvironment prior to sporozoite challenge. We conducted a time course of *P. chabaudi* BS infection, collecting blood and liver tissue at 10, 15, and 20 dpi, corresponding roughly to peak parasitemia, mid-resolution, and submicroscopic parasitemia, respectively (Fig. 1A). Multiple hallmarks of acute inflammation were visible in the liver during acute *P. chabaudi* BS infection, including hepatocyte necrosis and mononuclear cell infiltration, but were resolved by day 20 (Fig. 2A-B), with the exception of pigment-rich Kupffer cells, which we hypothesize are hemozoin-laden Kupffer cells. These cells increased gradually over the course of BS infection (Fig. 2C, fig. S2). There was no difference in serum levels of enzymatic markers of liver damage aspartate aminotransferase (AST) or alanine aminotransferase (ALT) levels, or their ratio, between uninfected and *P. chabaudi* experienced mice 20 dpi (Fig. 2D). Serum cytokine levels, including TNFa, IFNg, IL-6, and IL-10, each which has been previously associated with *Plasmodium* infection(*22, 23*), were also not significantly different from uninfected mice at 20 dpi (Fig. 2E). Taken together, these data are consistent with the resolution of liver pathology occurring concurrently with resolution of parasitemia in the *P. chabaudi* model as previously described (*24*).

**Fig. 2.**
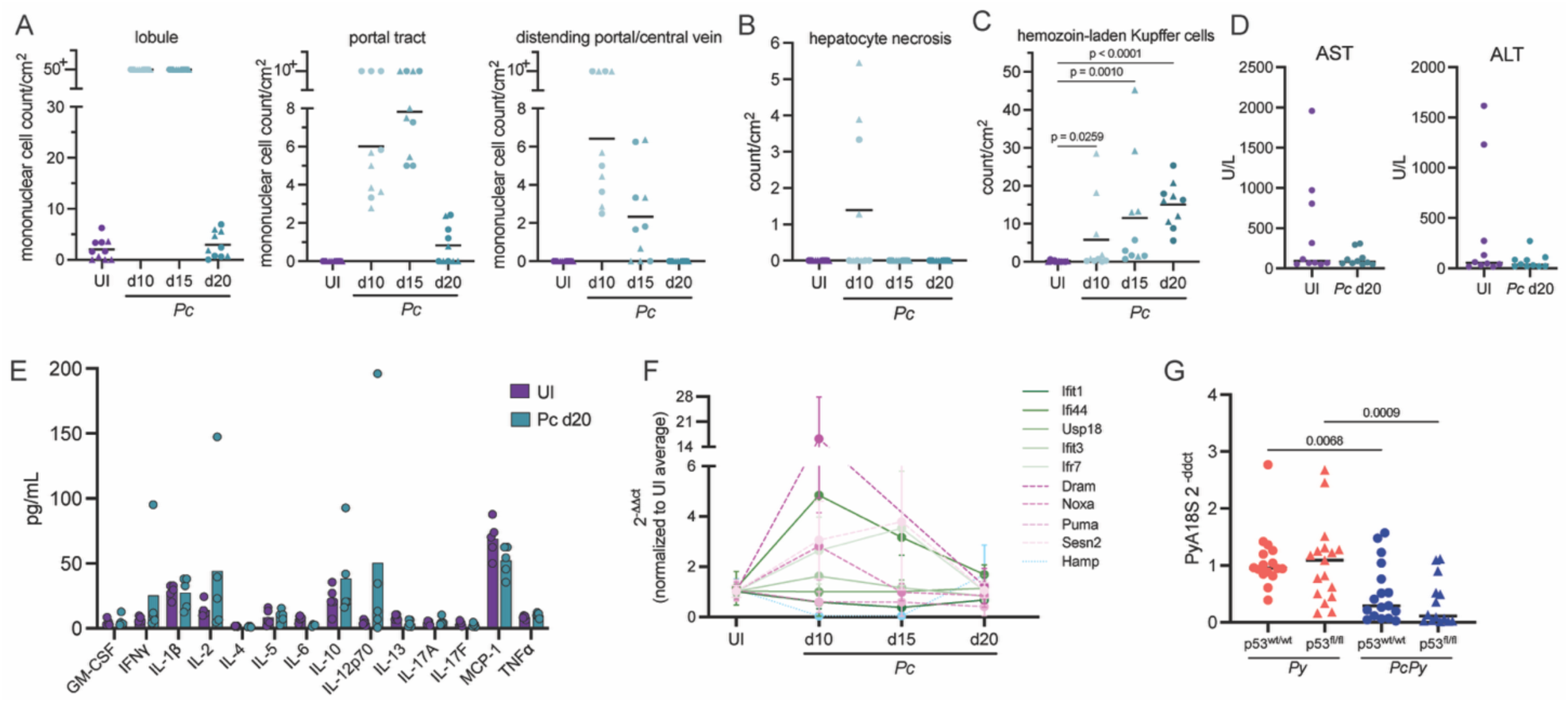
A majority of systemic inflammation and liver pathology are resolved at the time of LS challenge. **(A)** Area-normalized count of mononuclear cell foci in the liver lobule, portal tract, and distending portal or central vein over the time course of *P. chabaudi* BS infection. Each dot represents a single infected (Pc) or uninfected (UI) mouse (n = 5 male and 5 female mice). **(B)** Area-normalized count of foci of hepatocyte necrosis in the liver in UI and Pc mice. **(C)** Area-normalized count of pigment-rich (consistent with hemozoin-laden) Kupffer cells in the livers of UI or *Pc* mice over the time course. **(D)** Levels of AST and ALT measured in mouse serum 20 days post-*P. chabaudi* BS infection. **(E)** Serum cytokine levels in uninfected and day 20 *P. chabaudi*-infected mice. Each dot represents a single mouse. Data were analyzed by multiple Mann-Whitney tests. **(F)** Relative expression of hepcidin (*Hamp*), p53-regulated (*Noxa, Sesn2, Puma, Dram*), and IFN-stimulated (*Ifit1, Ifit3, Ifi44, Ifg7, Usp18*) genes measured by qPCR in the livers of female *Pc* BS-infected mice over a time course. Data were normalized to *Gapdh* and the average of UI mice. Data were analyzed by Dunnett’s multiple comparisons test. n = 5 female mice per time point. **(G)** PyA18S quantified by qPCR in livers of *Alb-cre*^+/+^*Trp53*^fl/fl^ or *Alb-cre*^+/+^*Trp53*^wt/wt^ mice 44h post-sporozoite infection. Mice were either naïve (*Py* only) or had reached submicroscopic *P. chabaudi* BS infection (*PcPy*) at the time of sporozoite challenge. Data were normalized to *Gapdh* and to the average of *Alb-cre*^+/+^*Trp53*^wt/wt^ *Py* only mice within each experiment. Each dot represents a single mouse from one of three experiments. Data were analyzed by Dunn’s multiple comparisons test.

Previous work has implicated hepcidin and interferon-related signaling in reducing LS infection that occurred during acute BS infection (*9, 11*). To assess this possibility, we measured expression levels of hepcidin (*Hamp*) and IFN-stimulated genes (*Ifit1, Ifit3, Ifi44, Ifr7, Usp18*) in livers over the time course of *P. chabaudi* BS infection. We also measured expression of a panel of p53-regulated genes (*Noxa, Sesn2, Puma, Dram*) within the liver, as suppression of p53 signaling has been shown to be crucial for successful *Plasmodium* LS infection in mouse models of infection (*25*) and increased p53 in circulating monocytic cells has been linked with protection from symptomatic infection in children (*26*). While expression of many of these target genes was elevated during acute BS infection, none remained significantly elevated 20 dpi (Fig. 2F). Elimination of the p53 gene in hepatocytes using an *Alb-cre*^+/+^*Trp53*^fl/fl^ floxed mouse model also did not rescue *P. yoelii* LS levels in *P. chabaudi* BS-experienced mice (Fig. 2G).

Since *P. chabaudi*-infected animals closely resembled uninfected animals in terms of inflammation at the time of *P. yoelii* sporozoite challenge, we reasoned that the suppression of liver infection might originate from a liver state that materialized after sporozoite challenge. To characterize the liver response to sporozoite challenge, we performed RNAseq on liver tissue collected over a time course post-*P. yoelii* sporozoite challenge from naïve and previously *P. chabaudi* BS-experienced mice (Fig. 3A-B, table S1). In the absence of LS infection, 468 genes were differentially expressed between *P. chabaudi* BS experienced and naïve animals (table S1). Gene ontology enrichment analysis revealed upregulation of largely metabolic biological processes (table S2), consistent with our previous data showing resolution of inflammation within the liver by 20 dpi (Fig. 2). Enrichment analysis of differentially expressed genes in naïve mice infected with *P. yoelii* sporozoites revealed few changes at 6 or 24 hpi and an upregulation of innate immune processes in the liver, including cytokine production and interferon responses, at 44 hpi, consistent with previous studies (Fig. 3C, Suppl table S2)(*27*). In *P. chabaudi* BS-experienced mice, we saw a similar enrichment of innate immune processes in response to *P. yoelii* LS. However, this signature in *P. chabaudi* experienced animals was evident by 24 hpi as opposed to 44 hpi in previously naïve animals (Fig. 3C-D, table S2). When comparing previously naïve with *P. chabaudi* BS-experienced mice at 44 hours post-*P. yoelii* LS infection we observed upregulated expression of genes enriched for interferon and virus responses (table S2), suggesting a subset of immune processes are induced earlier and to a greater extent in *P. chabaudi* BS-infected mice. While these inflammatory responses, when elicited at 44 hours close to the end of LS infection, can only modestly impact infection (*28*), an established response prior to the development of late LS infection could serve as a potent controller of LS burden.

**Fig. 3.**
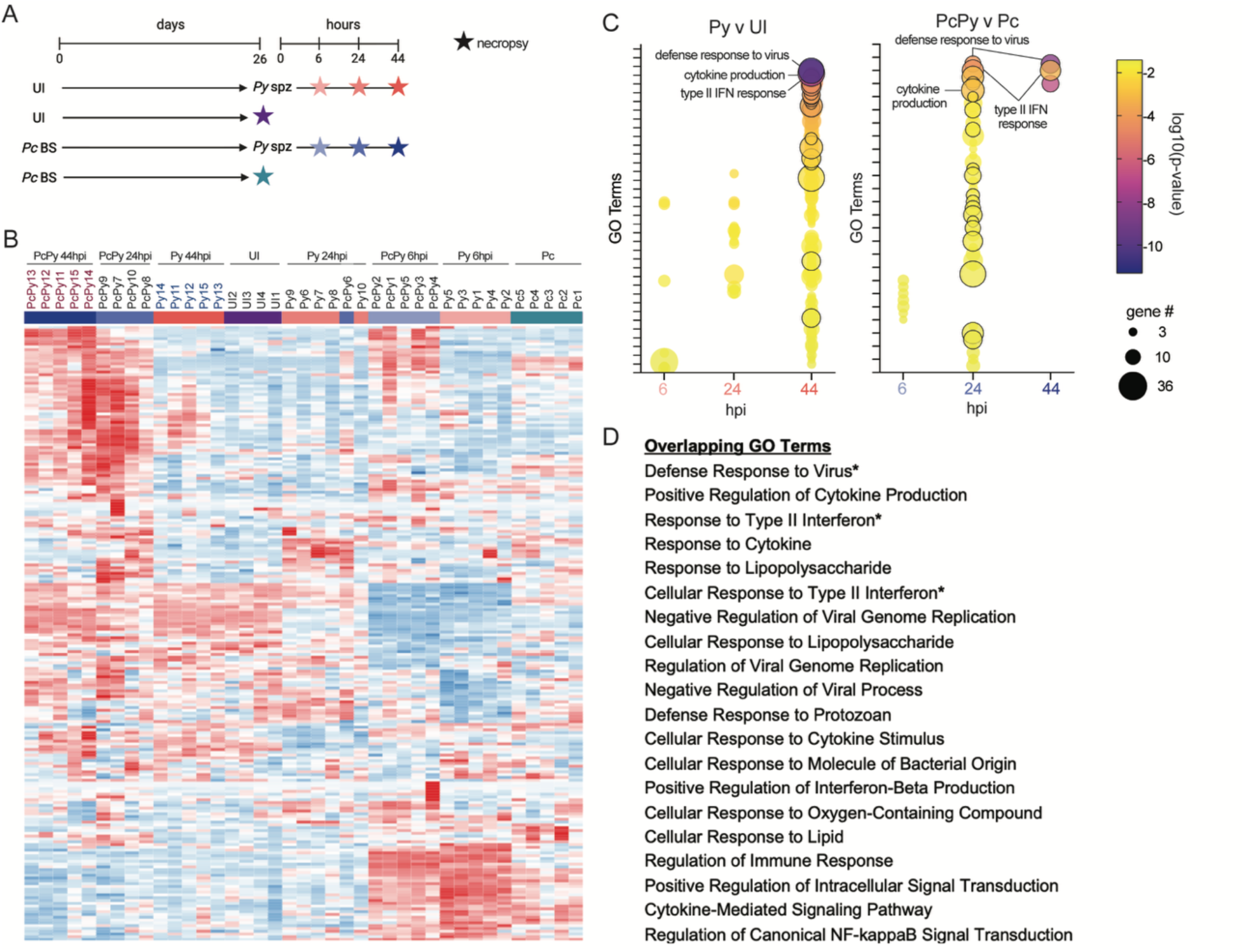
Mice with a prior *P. chabaudi* BS infection exhibit earlier upregulation of genes involved in immune processes within the liver upon *P. yoelii* LS infection compared to naïve mice. **(A)** Schematic of experimental timeline and treatment groups for RNAseq of whole liver tissue. Uninfected (UI) or *P. chabaudi* blood-stage (*Pc* BS) infected mice were either necropsied or infected with *P. yoelii* sporozoites (*Py* spz) at 26 dpi when *Pc* parasitemia was no longer detectable by microscopy. *Py* and *PcPy* mice were necropsied at 6, 24, and 44h post-*Py* spz infection (hpi). Stars indicate necropsies. **(B)** Clustergrammer heat map showing normalized gene expression values across all samples. Data were normalized to logCPM. Each column represents an individual mouse. **(C)** Bubble plots showing Gene Ontology (GO) enrichment analysis of genes with upregulated expression in *Py* (normalized to UI) and *PcPy* (normalized to *Pc*) mice at 6, 24, and 44 hours post-*Py* infection. FDR-adjusted p-value < 0.05 was used as a cut-off. Overlapping GO terms enriched in *Py* and *PcPy* groups at any time point are outlined in black. The top three terms significantly enriched in *Py* v UI are labeled. **(D)** Overlapping GO terms enriched in *Py* v UI and *PcPy* v *Pc* ordered by p-value in *Py* v UI. All terms were enriched 44 hpi in *Py* and 24 hpi in *PcPy*. Asterisks indicate terms also enriched at 44 hpi in *PcPy*.

Innate memory is characterized by an altered transcriptional response, either repressed (tolerance) or enhanced (trained immunity), that occurs in an antigen-independent manner and is driven by epigenetic changes (*29*). We hypothesized that *P. chabaudi* BS infection induces a trained immune state within the liver (Fig. 4A), as indicated by the more rapid and robust induction of immune gene expression upon sporozoite infection (Fig. 3C-D). To interrogate the epigenetic state of the liver we performed ATAC-seq on whole liver tissue from uninfected and *P. chabaudi* BS-experienced mice at 20 dpi. A multitude of genes exhibited differential accessibility, suggesting that BS infection has the capacity to alter the epigenetic landscape of liver cells that is retained after the resolution of acute parasitemia (Fig. 4B, table S2). We observed that 792 genes had increased, and 317 decreased, accessibility compared to naïve controls, corresponding to 6.3% and 2.5% of genes with mapped fragments, respectively (adjusted p-value < 0.05 and fold-change > 1.5 or < 0.75) (table S2). Gene ontology enrichment analysis revealed increased accessibility of genes involved in immune system processes and decreased accessibility of genes with a role in various metabolic and catabolic processes (Table S3). We then asked what genes exhibited both increased accessibility *and* increased transcription in *P. chabaudi* experienced mice at 20 dpi upon *P. yoelii LS* challenge as compared to previously naïve and *P. chabaudi* only controls (Fig. 4B-C). We identified a group of interferon-stimulated genes (ISGs), Gm12250, Gbp4, Gbp8, Gbp9, Iigp1, and Ifi47, with earlier and more pronounced peaks in expression levels upon *P. yoelii* LS infection in *P. chabaudi*-trained mice and increased chromatin accessibility (Fig. 4C-D). All of these genes are interferon-inducible GTPases, a family of proteins involved in regulating infection of multiple pathogens, including *Plasmodium* and other apicomplexan parasites (*30–33*), including during LS infection. Interestingly, we also observed increased accessibility and increased expression of a group of major histocompatibility complex (MHC) II genes which can be aberrantly expressed by hepatocytes in response to IFN stimulation and cause the cell to act as a non-professional antigen presenting cell (*34*). It is tempting to speculate that this reprogrammed hepatocyte state in BS experienced animals might also rewire how the liver responds to T cell responses. Previous work has demonstrated that a shift in metabolic activity is a major driver of innate memory training. Specifically, trained cells have elevated glycolytic activity, which was attributed to increased accessibility and expression of glycolytic genes such as 6-phosphofructo-2-kinase/fructose-2,6-biphosphatase 3 (PFKFB3)(*35*). Both of our RNA-seq and ATAC-seq data revealed increased accessibility and transcription of the PFKFB3 locus between Pc-experienced and uninfected mice (table S1, table S3), suggesting that a rewired metabolic state in the liver could underly immune training.

**Fig. 4.**
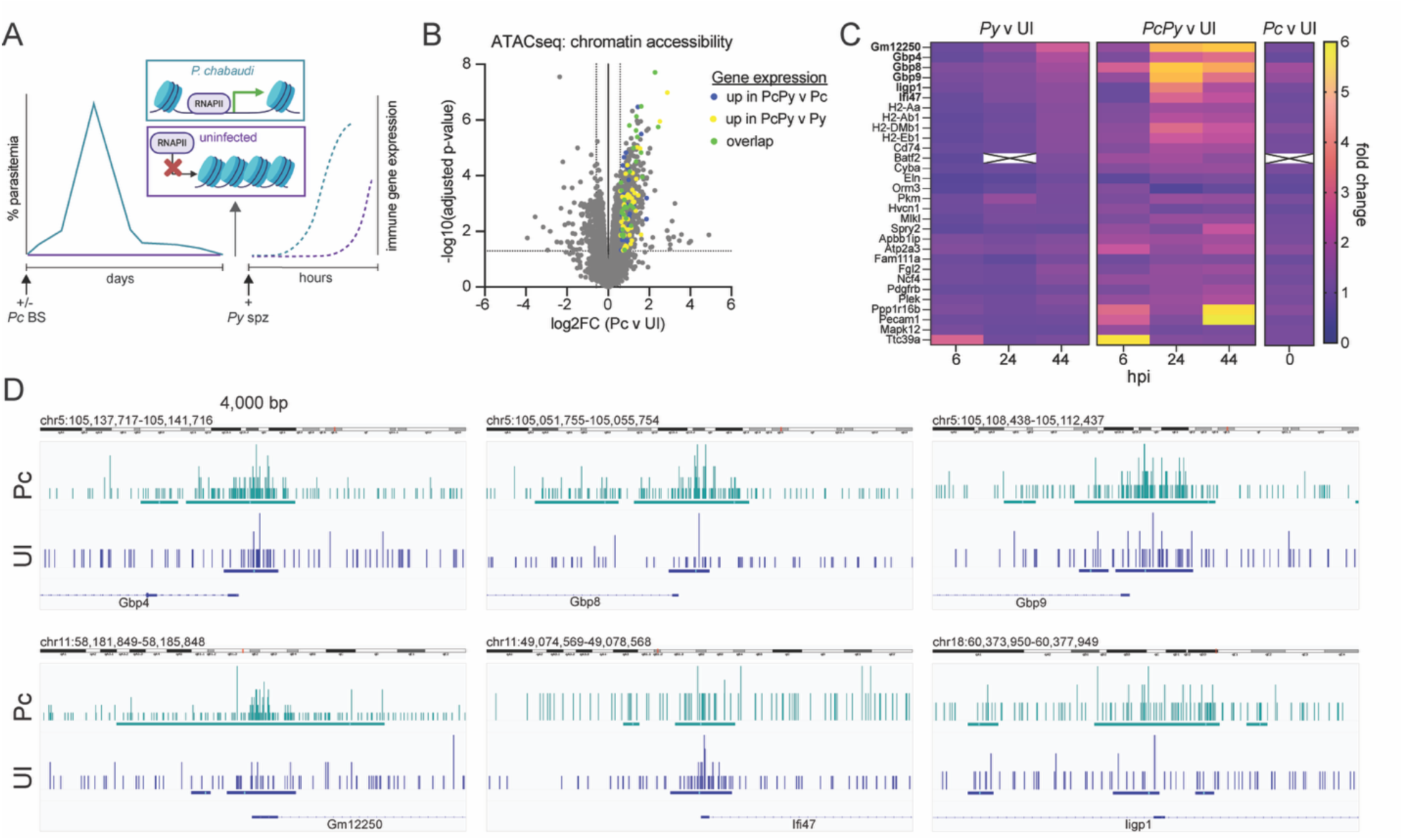
Overlap in expression kinetics and chromatin accessibility suggests epigenetic changes induced by *P. chabaudi* BS infection may underlie an enhanced IFN response within the liver upon *P. yoelii* sporozoite infection. **(A)** Hypothesis schematic: *P. chabaudi* blood-stage infection leads to changes in chromatin accessibility, which allow for more rapid and robust immune gene expression in the liver upon *P. yoelii* sporozoite challenge. **(B)** Differential chromatin accessibility between d20 *Pc* and UI mice measured by ATACseq. Each dot represents a gene. Fold change > 1.5 and adjusted p-value < 0.05, indicated by dotted lines, were used as cut-offs for significance. Genes with increased accessibility in *Pc* v UI mice that also exhibited increased expression in *PcPy* v Pc mice (blue), *PcPy* v *Py* mice (time-matched) (yellow), or both (green), at any time point post-infection, are indicated. **(C)** Heatmap showing expression levels of subset of genes with increased chromatin accessibility and increased expression in *PcPy* v *Pc* and *PcPy* v *Py* mice. Data are shown as mean fold change normalized to UI mice. Interferon-stimulated genes are indicated in bold. X indicates expression below detectable levels. **(D)** Representative ATACseq tracks showing increased chromatin accessibility at IFN-responsive genes. Created using Integrative Genomics Viewer (*50*).

Trained immunity has been described in circulating innate immune cells and in hemopoietic stem cells in the context of *Plasmodium* infection (reviewed in (*13*)) and in tissue-resident macrophages, including Kupffer cells, in other contexts (*36, 37*). Epigenetic changes in hepatocytes have been documented (*36*) and a trained immunity phenotype described in liver cells in a Zebrafish larva model (*38*), however, the capacity for mammalian hepatocytes to undergo innate memory has not been described to date. Hepatocytes are the predominant cell type in the liver and are heavily polyploid in mice (*39*), however it remains possible that a sufficiently robust change in gene expression in another cell type, such as Kupffer cells, could be responsible for the phenotype we see in whole liver tissue. To determine whether hepatocytes respond more rapidly to secondary challenge after BS experience, we isolated hepatocytes from uninfected and *P. chabaudi* BS experienced mice 20 dpi and plated these cells in culture for stimulation *ex vivo* (Fig. 5A)(*40, 41*). Having training occur *in vivo* allowed the maintenance of any contributions of other cell types or the tissue microenvironment to the establishment of innate memory (*42, 43*), while secondary stimulation *ex vivo* allowed us to distinguish a true recall response from any prolonged effects of hemozoin retained in Kupffer cells, which is consistent with the pigment-rich cells we observe (Fig. 2, fig. S2C), and has been implicated in modulation of immune functions (*44*). In purified hepatocytes we observed significantly reduced expression of Cd45 and Clec4f, expressed by leukocytes and Kupffer cells, respectively, compared with the nonparenchymal and pre-purification cell populations (fig. S3A), suggesting we isolated a highly hepatocyte-enriched population. To assess the recall capacity of these hepatocytes, we stimulated cells with a molecule not derived from *Plasmodium* antigens, double-stranded calf thymus DNA (ctDNA), which has been shown to robustly induce an ISG response similar to that of *Plasmodium* infection (*45*). We monitored expression of the panel of ISGs (Fig. 4C) in isolated hepatocytes over a time course from 0 to 8 hours post-stimulation via qPCR. Hepatocytes isolated from *P. chabaudi* BS infected mice 20 dpi exhibited more rapid induction of all six of the genes in the ISG panel in response to treatment when compared to hepatocytes isolated from naïve mice (Fig. 5B). Hepatocytes from *P. chabaudi* infected mice did not exhibit increased ISG expression at baseline compared to those from naïve mice, in fact some genes exhibited reduced levels pre-stimulation, (fig. S3B) indicating the enhanced response is not a residual effect of *P. chabaudi* infection, but rather an accelerated and enhanced recall response to a secondary stimulus.

**Fig. 5.**
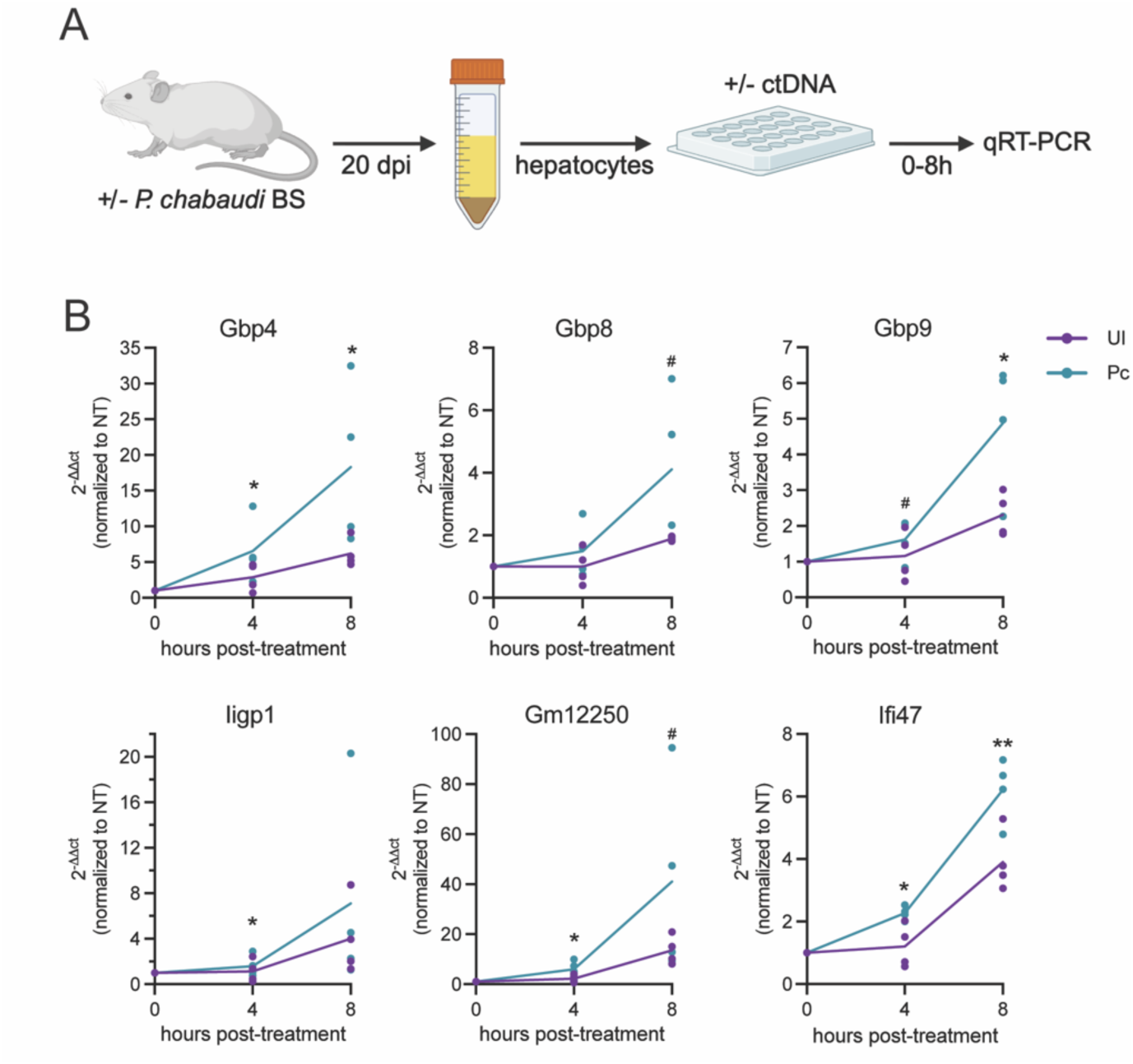
Hepatocytes exhibit trained immunity in response to blood-stage infection. **(A)** Experimental schematic: Livers were perfused and hepatocytes isolated from *P. chabaudi* blood-stage (BS)-infected mice after reaching submicroscopic parasitemia (20-26 dpi) or from uninfected age-matched controls. Plated hepatocytes were stimulated with or without 2.5μg/mL ctDNA, and cells collected in TRIzol over a time course for qPCR. **(B)** IFN-stimulated gene expression in hepatocytes isolated from UI and Pc infected mice and stimulated *ex vivo* with ctDNA. Data were normalized to Gapdh and to unstimulated time- and infection status (UI or Pc) matched hepatocytes. Each dot represents a single biological replicate conducted with cells pooled from 1 to 3 mice. Data were analyzed by a ratio paired t-test.

Studies of innate immune memory have largely been focused on innate immune cells and, more recently, stem and endothelial cells (*46, 47*). Here we give, to our knowledge, the first report that mammalian hepatocytes can also be trained. Specifically, *Plasmodium* BS infection epigenetically reprograms hepatocytes to more rapidly respond to secondary immune stimulation in an antigen-independent manner. Hepatocytes are long-lived cells with an average age of almost 3 years in an adult liver (*48*), allowing time for innate immune memory to impact sequential infections in the field. While the focus of this study is on *Plasmodium* infection, hepatocytes harbor a wide variety of pathogens and have numerous functions that impact mammalian biology, including the detoxification and processing of drugs (*49*). Further work will determine the extent of the impact trained immunity in hepatocytes: whether it can be induced by other stimuli and if its biological impact extends beyond regulating *Plasmodium* liver infection.

## Acknowledgements

Funding

National Institutes of Health grant R01AI158719 (AK, NWS, TMT)

National Institutes of Health grant R01AI177257 (AK, EKKG, NKM, TT)

## Author contributions

Conceptualization: AK, EKKG, NH, NWS, TMT

Investigation: EKKG, TT, VIP, NS, LW, LV, CF, BV, AS, RPK

Supervision: SM, NKM, AK

Writing – original draft: EKKG

Writing – review & editing: AK, EKKG, NH, RBP, RPK, NKM, NWS, TMT

## Data, code, and materials availability

RNAseq and ATACseq data are deposited on GEO, accession numbers GSE338810 and GSE338812, respectively. All other data are available in the main text or the supplementary materials.

## Supplementary Materials for Materials and Methods

### Parasite production

Swiss Webster mice (Envigo) were used to produce *Plasmodium yoelii* 17XNL and *Plasmodium chabaudi* parasites. *P. yoelii* sporozoites were obtained by infecting female Swiss Webster mice with BS *P. yoelii* parasites. Upon observation of gametocyte exflagellation, these mice were used to feed *Anopheles stephensi* mosquitoes. Fourteen to fifteen days post-blood meal, salivary glands were dissected, and sporozoites were isolated by manual grinding in cold Schneider’s solution, followed by centrifugation at 1000rpm for 2 minutes. The supernatant containing sporozoites was collected and quantified using a hemocytometer. For *P. chabaudi* BS infections, cryopreserved *P. chabaudi*-infected red blood cells (RBCs) (10% parasitemia stock, 1:1 in freeze media consisting of 28% glycerol, 3% sorbitol, and 0.65% NaCl in Alsever’s solution) were thawed and injected intraperitoneally (IP) into a Swiss Webster mouse (Envigo) to generate a starter infection. Once parasitemia reached 1%, blood was collected via the tail vein and used for IP infection of experimental mice.

### Mouse infections

All experimental infections were performed using 8–10-week-old female C57BL/6J mice (Jackson Laboratories), unless indicated otherwise. For conditional hepatocyte-specific p53 knockout experiments, Alb-cre mice (B6. Cg-Speer6-ps1^Tg(Alb-cre)21Mgn^/J) were crossed with p53^LoxP^ (B6.129P2-Trp53^tm1Brn^/J) mice (Jackson laboratories). F1 heterozygous progeny were intercrossed, and offspring were genotyped using a standard PCR protocol (Jackson Genotyping Protocol #23419). Male and female homozygous *Alb-cre*^+/+^*Trp53*^fl/fl^ and *Alb-cre*^+/+^*Trp53*^wt/wt^ mice were then used for experimental infections. All animal procedures were conducted in accordance with protocols approved by the Seattle Children’s Institutional Animal Care and Use Committee.

Mice were first infected at 8-10 weeks with *P. chabaudi* via IP injection of 1x10^5^ infected RBCs diluted in PBS. Parasitemia was monitored every other day by Giemsa-stained thin blood smears starting 4 days post-infection (dpi) until parasites were no longer detectable by microscopy (10 fields of view under 100x oil immersion). Once *P. chabaudi* parasitemia fell below the detection threshold (20–25 dpi), mice were challenged with 25,000 *P. yoelii* 17XNL sporozoites by retro-orbital injection. At the time of *P. yoelii* infection, blood spots were collected on Cytiva Whatman 903 Protein Saver Cards (Fisher Scientific) for quantification of *P. chabaudi* 18S rRNA via qRT-PCR. Mice were euthanized 44 hours after *P. yoelii* challenge. Blood was collected via cardiac puncture. Blood samples were allowed to clot at room temperature for at least 30 minutes. Serum was isolated by centrifugation at 1,500 g for 15 minutes at 4°C and stored at −80°C. Livers were collected and portions flash-frozen in liquid nitrogen, fixed in 4% paraformaldehyde (PFA) for histological analysis, or homogenized in TRIzol reagent (Thermo Fisher) for RNA extraction.

### CD4/8 depletion and flow cytometry

After *P. chabaudi* parasitemia dropped below detectable levels, mice were intraperitoneally injected with 400 µg of anti-CD4 (clone GK1.5, SelleckChem) or anti-CD8 (clone 2.43, gift from Dr. Nathan Schmidt) monoclonal antibodies on days −3 and −1 prior to *P. yoelii* sporozoite challenge. Depletion efficiency was assessed by flow cytometry on peripheral blood collected at the time of sporozoite infection. A spleen from an uninfected mouse was used for single antibody stain compensation controls. The spleen was mechanically dissociated through a 100 µm strainer into flow buffer (PBS + 2% FBS). Blood was collected in heparinized tubes. Red blood cells were lysed with ACK buffer for 15 minutes at room temperature, then cells were washed, counted, and resuspended in flow buffer.

Two million splenocytes or all available blood leukocytes were plated in v-bottom 96-well plates. Fc receptors were blocked with anti-mouse CD16/32 (1:100, Biolegend) for 15 minutes at 4°C. Cells were stained with the following antibodies (Biolegend): CD3-PE, CD4-BV750, and CD8-FITC, and Aqua Zombie live/dead stain (BioLegend) for 30 minutes at 4°C. All antibodies were used at a 1:100 dilution. Cells were fixed in 4.2% paraformaldehyde for 15 minutes at 4°C and washed twice in flow buffer. Samples were run on a BD Symphony A5 flow cytometer and analyzed with FlowJo software (FlowJo_v10.10.0, BD Biosciences). Compensation was performed using single-stained cell samples prepared from uninfected spleen tissue. Lymphocytes were gated based on FSC/SSC profile, with singlets and live (Aqua Zombie-negative) cells selected for further analysis (fig. S1G). T cells were identified as CD3⁺, and CD4⁺ and CD8⁺ subsets were quantified as a percentage of CD3⁺ cells.

### qRT-PCR

Total RNA was extracted from homogenized liver tissue using TRIzol (Thermo Fisher Scientific) and chloroform. cDNA was synthesized using the iScript gDNA Clear cDNA Synthesis Kit (Bio-Rad) according to the manufacturer’s instructions. Quantitative real-time PCR (qRT-PCR) was performed using SYBR Green on a QuantStudio 5 system (Applied Biosystems) using the primer sequences in Table S4. All reactions were run in triplicate. Gene expression was normalized to *Gapdh* expression and analyzed using the ΔΔCt method (*51*). For parasite quantification, *PyA18S* was used to assess relative *P. yoelii* LS burden. Data were normalized to the average ΔCt of the control group within each experimental replicate.

### Blood spot quantification

Dried blood spot (DBS) samples collected on Whatman® Protein Saver 903 cards (Cytiva; Cat. #10534612) were manually excised using sterile scissors and transferred into nuclease-free tubes. Each sample was then lysed with 2 mL of NucliSens® Lysis Buffer (bioMérieux, Inc., Durham, NC, USA; Cat. #280134) and incubated at 55°C for 30 minutes with agitation every 10 minutes. Total nucleic acids were extracted using the NucliSens® easyMAG® system (bioMérieux, Inc.) by loading 1 mL of lysed sample per extraction. The eluates were subjected to quantitative reverse transcription PCR (qRT-PCR) using the SensiFAST™ Probe Lo-ROX Kit (Bioline, London, UK; Cat. #BIO-78005) on a QuantStudio™ 5 Real-Time PCR System (Thermo Fisher Scientific, Waltham, MA, USA) under conditions described previously (*52*). Amplification was done using PCR primers and probes targeting mouse GAPDH (IDT Inc., Coralville, IA, USA) and Pan-*Plasmodium* 18S rRNA as described (*53*). *Plasmodium* 18S rRNA copy numbers were quantified using a custom-manufactured lot of Armored RNA (Asuragen, Austin, TX, USA) containing full-length *Plasmodium* 18S rRNA. Standard curves were generated from serial dilutions of this quantified RNA and used to interpolate copy numbers in study samples, as described previously (*52, 54, 55*).

### Histology and pathology scoring

After fixation in 4% PFA for 24 hours, liver tissue was paraffin-embedded, and 4mm sections taken for staining by the Microscopy and Histopathology CoLab at Seattle Children’s Research Institute. From each mouse, a hematoxylin and eosin-stained formalin-fixed paraffin-embedded section of liver ranging between 35 and 180 mm^2^ (mean +/- SD, 95.5 +/- 41.2 mm^2^) was examined histologically. For each of the following histological features, the number of discrete sites per section was quantified and used to determine the relative abundance of each finding per cross-sectional area: lobular inflammation (+/- hepatocyte necrosis), portal inflammation, portal or hepatic venous distension by inflammatory cells, and confluent hepatocyte necrosis without inflammation. Prussian blue stain was used to detect iron deposition. Pigment-rich cells were quantified in tissue sections stained with hematoxylin and diaminobenzidine using QuPath Image Analysis software.

### RNA-seq

Total RNA was extracted using TRIzol and chloroform and quality assessed using an Agilent 5400 Fragment Analyzer. One sample was excluded using a RIN cutoff of 5. The remaining samples were sequenced by Novogene using paired-end sequencing using the NovaSeq X Plus platform. Sequences were aligned and differential gene expression and enrichment analysis performed using the BioJupies platform (https://maayanlab.cloud/biojupies/) (*5C*) with default parameters. RNA-seq FASTQ files are available on GEO under accession number GSE338810.

### ATAC-seq

Perfused liver tissue was collected at necropsy and flash frozen in liquid nitrogen. Nuclei isolation and transposition were conducted according to the protocol developed by Corces et al. for frozen tissue (*57*). ATAC resuspension buffer (RSB), RSB-tween (RSB-T), and homogenization buffer unstable solution (HBUS) were all made as previously described (*57*). For each sample 10-20mg of tissue was homogenized in HBUS using a Dounce, with 10 strokes of pestle A, followed by 20 strokes of pestle B, and then filtered through a 70mm Flowmi strainer (Fisher Scientific). Samples were pelleted by centrifugation at 350g for 5 minutes at 4°C and resuspended in HBUS. Nuclei were isolated using a 25-30-40% iodixanol gradient spinning at 3000g for 20 minutes in a swinging bucket centrifuge with the brake off. Nuclei were quantified using trypan blue staining and diluted in RSB-T to obtain 50,000 nuclei in 1mL for each sample. Nuclei were then pelleted at 500g for 10 minutes and resuspended in 50 μL transposition mix (25 μL 2x TD buffer, 2.5 μL TDE1 enzyme (Illumina), 16.5μL PBS, 0.5 μL1% digitonin, 0.5 μL 10% Tween-20, 5 μL water). The transposition reaction was carried out in a thermomixer at 1000 rpm, 37°C for 30 minutes, then processed using a clean and concentrator-5 kit (Zymogen). The remainder of the library preparation was performed as described by Buenrostro et al. (*58*). Purified transposed DNA was amplified for 5 PCR cycles using NEBNext High-Fidelity 2X PCR Master Mix (New England Biolabs) and a unique adaptor primer combination for each sample (Table S5). Samples were then put on ice, and qPCR was performed using 5 μL of each sample to determine the number of additional cycles needed to amplify samples while avoiding saturation (cycle number corresponding to 1/3 of maximum fluorescent intensity). Fully amplified samples were purified using the Zymogen Clean and concentrator-5 kit and quantified using the Qubit dsDNA HS Assay (ThermoFisher) for pooling.

The pooled library was sequenced by Psomagen using the Illumina NovaSeq 6000 platform for 151-bp paired-end reads. ATAC-seq FASTQ files are available on GEO under accession GSE338812. Data QC, alignment, and peak calling were conducted using the PEPATAC pipeline as described (*59*). Pre-alignment was performed by aligning to the mouse mitochondrial genome before the main alignment to the primary reference genome. Both *Mus musculus* mitochondrial genome (mouse_chrM2x) and primary reference genome (mm10) assets were downloaded using refgenie (https://refgenie.databio.org/en/latest/)(C0). Transcription start site (TSS) annotation for mm10 genome assembly was downloaded from the ENCODE portal (https://www.encodeproject.org, accession ENCFF498BEJ). The genomic region blacklist was obtained from the ENCODE blacklist repository (https://github.com/Boyle-Lab/Blacklist)(C1). Quality control was done as part of the PEPATAC pipeline using the default settings. Peak annotation was performed using HOMER version 5.1 (http://homer.ucsd.edu/homer/)(C2). Differential accessibility analysis was performed using edgeR (https://bioconductor.org/packages/release/bioc/html/edgeR.html, version 4.6.3)(*63*). Enrichment analysis was performed using the DAVID functional annotation tool with the set of 12,487 identified genes as background.

### Liver perfusion and *ex vivo* hepatocyte stimulation

Hepatocytes were isolated from age-matched female c57bl/6 mice that were uninfected or had reached submicroscopic parasitemia after infection with *P. chabaudi* BS parasites (21-26 dpi) based on previously published protocols (*40, 41*). Livers were perfused with 15mL of warmed DBPS (Ca/Mg-Free) containing 500 µM EGTA, followed by 50mL of DBPS containing 1mM CaCl2 and 0.1mg/mL Liberase TL (Roche). The liver was then removed and gently pulled apart in hepatocyte culture media (HCM): DMEM containing 10% FBS and 1% penicillin-streptomycin. The cellular suspension was then filtered through a 100 µm cell strainer, followed by a 70 µm cell strainer into fresh media. Cells were then spun at 50g, and the pellet resuspended in a 33% percoll solution. Cells were spun at 1000g for 3 minutes, and the pelleted hepatocytes resuspended in HCM. A final spin was done at 600g, and cells resuspended in HCM and seeded in collagen-coated 24-well plates at 1x10^5^ cells per well. Cells were allowed to adhere overnight before stimulation. Aliquots of total liver cells (post-filtration), non-parenchymal cells (supernatant after 50g spin), and purified hepatocytes were collected in TRIzol for RNA extraction.

Media was replaced and cells stimulated with 2.5μg/mL ctDNA (Thermo Fisher) and 3μg/mL lipofectamine 2000 (Invitrogen) or a diluent control. Supernatant was collected and stored at -80 °C and cells were collected in TRIzol for RNA extraction at 0, 4, and 8 hours post-treatment. RNA was extracted and qPCR run as described above. Data were normalized to Gapdh, and then to source (Pc v UI) and time-matched untreated controls. IFN levels were measured in supernatant by Eve Technologies using a Luminex 200 platform.

**Fig. S1.**
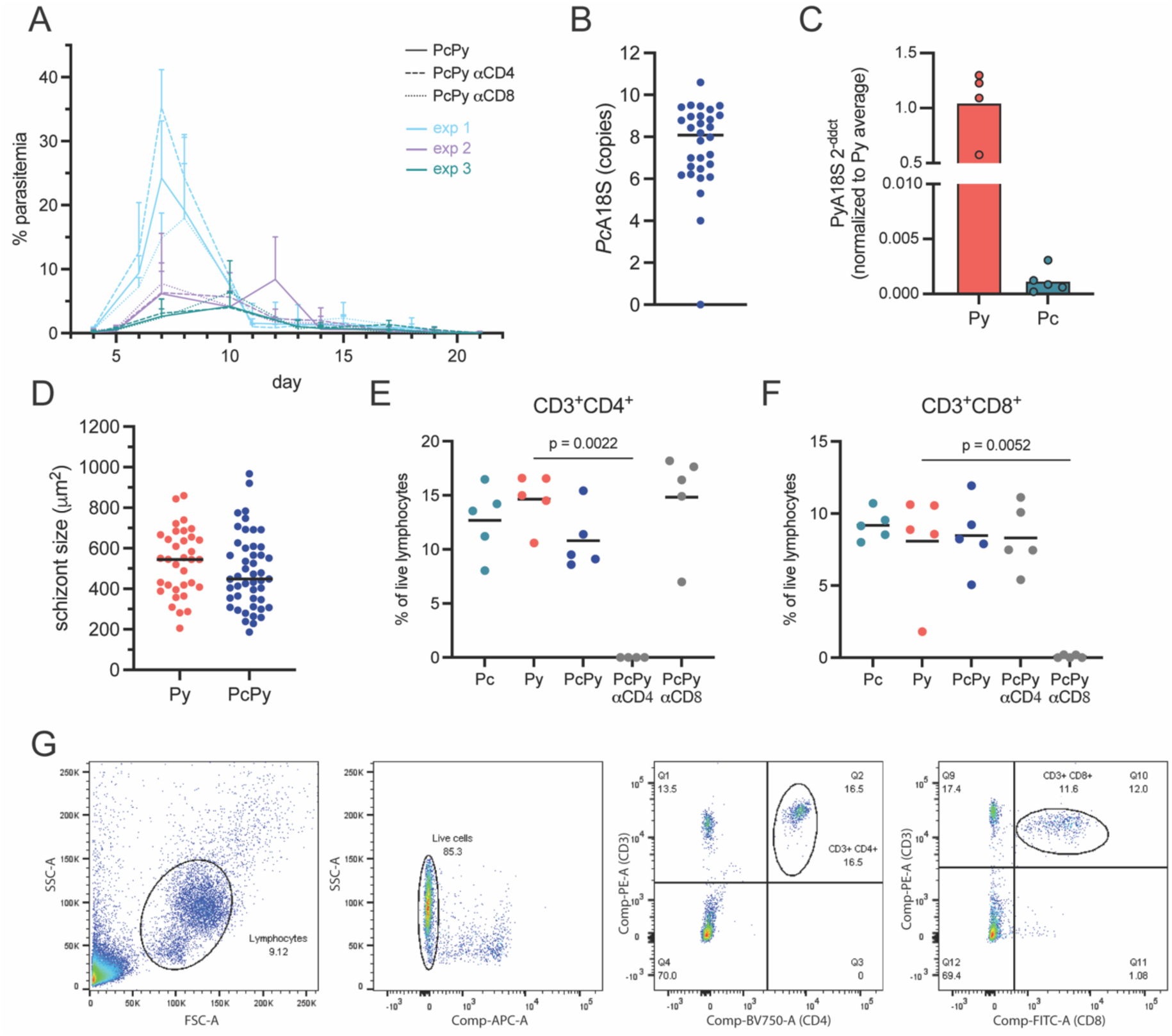
(A) *P. chabaudi* parasitemia shown as % infected red blood cells over time for each of 3 experimental replicates. Each replicate is indicated by a different color. Data are shown as mean and standard deviation of 5 mice per group per experiment. **(B)** PcA18S copy number detected in blood spots collected at the time of sporozoite challenge. Each dot represents a mouse from one of 2 independent experiments. **(C)** PyA18S levels were measured by qPCR in the livers of mice 44h after infection with *P. yoelii* sporozoites (*Py*) or 20 days post-infection with *P. chabaudi* blood-stage parasites (*Pc*). Each dot represents a single mouse. **(D)** *P. yoelii* schizont size measured in HCE-stained tissue 44h post-infection in previously naïve (Py) or day-20 *P. chabaudi*-infected (*PcPy*) mice. Each dot represents a single parasite, with 5 mice per group. Quantification of **(E)** CD3^+^CD4^+^ and **(F)** CD3^+^CD8^+^ live lymphocytes from a representative experiment. Each dot represents a single mouse. Data were analyzed by ANOVA with Dunn’s multiple comparisons test. **(G)** Representative gating strategy for measuring CD4^+^ and CD8^+^ T cell depletion in blood. Lymphocytes were gated by size, negative live/dead staining, positive CD3 staining, and positive CD4 and CD8 staining.

**Fig. S2.**
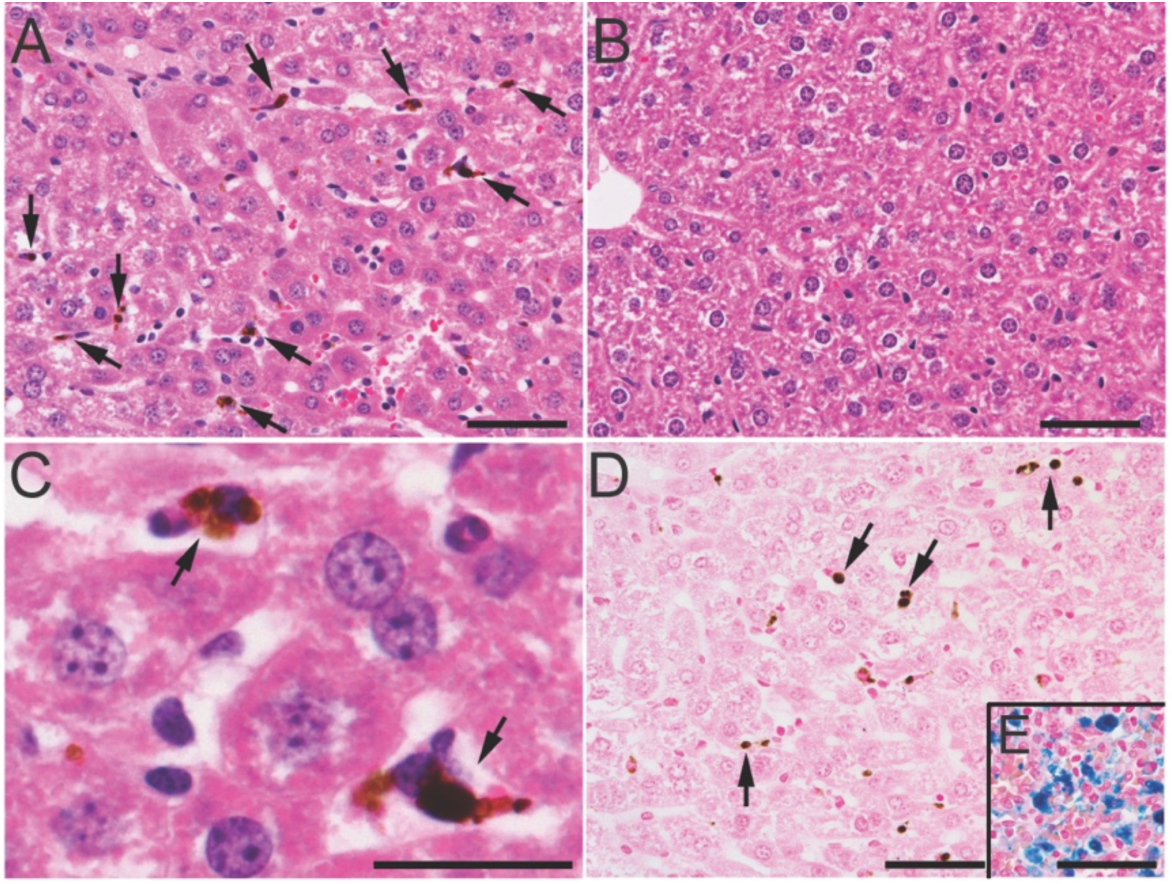
(A) Representative image of pigment-rich (consistent with hemozoin-laden) Kupffer cells in a *P. chabaudi-*infected liver 20 dpi with HCE staining. **(B)** Representative image of an HCE-stained liver from an age- and sex-matched uninfected mouse. **(C)** A higher magnification image of pigment-rich deposition is shown in (A). **(D)** Representative images of negative iron stain in d20 Pc tissue and **(E)** a positive control. Black arrows indicate areas of dense pigment, likely hemozoin deposition in Kupffer cells. Scale bars are 100μm for A, B, D, E, and 25 μm for C.

**Fig. S3.**
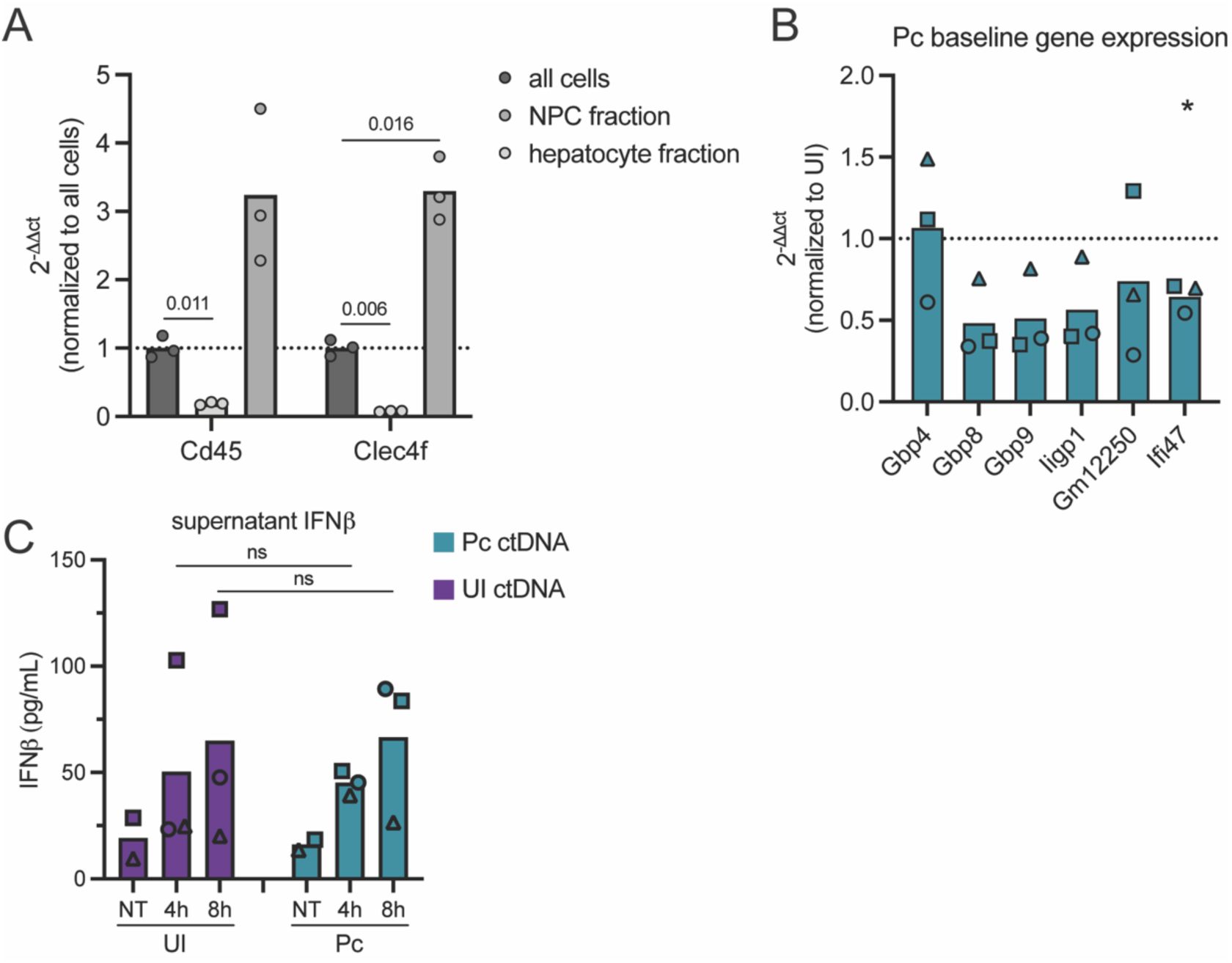
(A) Quantification of Cd45 and Clec4f expression by qPCR in cells isolated from mouse livers. Livers were perfused and aliquots of cells were taken after filtering (all cells), from the supernatant after centrifugation (non-parenchymal cells (NPCs)), and from the pellet after centrifugation in percoll (hepatocytes). Data were normalized to Gapdh and to the “all cells” sample. Each dot represents an aliquot. Data were analyzed by multiple unpaired t-test. **(B)** Expression levels of interferon-responsive genes in hepatocytes isolated from a *P. chabaudi* infected mouse 20 dpi at baseline (before stimulus with ctDNA *ex vivo*). Data were normalized to Gapdh and to matched samples from an uninfected mouse, set at 1. Each point represents a replicate with matched samples indicated by shape. Data were analyzed by one sample t-test. **(C)** IFNβ levels in hepatocyte supernatant at indicated time points post-ctDNA treatment *ex vivo*.

### Table S1. (separate file)

**RNAseq differential gene expression.** Differential gene expression analysis of RNAseq data from the livers of mice that were uninfected (UI), 20 days post-*P. chabaudi* BS (*Pc*), singly infected with *P. yoelii* LS (*Py*), or sequentially infected with *P. chabaudi* BS followed by *P. yoelii* LS (*PcPy*). Livers were collected at 6, 24, and 44 hours post-*P. yoelii* sporozoite challenge. Differential analysis was conducted between **(A)** *Py* v UI, **(B)** PcPy v Pc, **(C)** PcPy v Py, **(D)** *Pc* v UI, and **(E)** *PcPy* v UI groups. Genes are sorted by False Discovery Rate (FDR) adjusted p-value.

### Table S2. (separate file)

**RNAseq gene ontology enrichment analysis.** Gene ontology biological process enrichment analysis of genes with significantly upregulated and downregulated expression in Suppl Table 1. Adjusted p-value < 0.05 was used as a cut-off.

### Table S3. (separate file)

**ATACseq differential gene accessibility. (A)** Differential gene accessibility and **(B)** gene ontology biological process enrichment analysis for *P. chabaudi* infected and uninfected female mice 20 dpi determined by ATACseq. Adjusted p-value < 0.05 was used as a cut-off.

### Table S4. (separate file)

Primer sequences used for qPCR.

### Table S5. (separate file)

Adaptor primer sequences used for ATACseq.

## References

1. M. G. Netea et al., Defining trained immunity and its role in health and disease. Nat Rev Immunol 20, 375–388 (2020).

2. M. G. Netea, L. A. B. Joosten, Trained innate immunity: Concept, nomenclature, and future perspectives. J Allergy Clin Immunol 154, 1079–1084 (2024).

3. A. Dagenais, C. Villalba-Guerrero, M. Olivier, Trained immunity: A "new" weapon in the fight against infectious diseases. Front Immunol 14, 1147476 (2023).

4. T. M. Tran et al., An intensive longitudinal cohort study of Malian children and adults reveals no evidence of acquired immunity to Plasmodium falciparum infection. Clin Infect Dis 57, 40–47 (2013).

5. P. D. Crompton et al., Malaria immunity in man and mosquito: insights into unsolved mysteries of a deadly infectious disease. Annu Rev Immunol 32, 157–187 (2014).

6. I. Rodriguez-Barraquer et al., Quantification of anti-parasite and anti-disease immunity to malaria as a function of age and exposure. Elife 7, (2018).

7. L. C. Okell, A. C. Ghani, E. Lyons, C. J. Drakeley, Submicroscopic infection in Plasmodium falciparum-endemic populations: a systematic review and meta-analysis. J Infect Dis 200, 1509–1517 (2009).

8. C. Whittaker et al., Global patterns of submicroscopic Plasmodium falciparum malaria infection: insights from a systematic review and meta-analysis of population surveys. Lancet Microbe 2, e366–e374 (2021).

9. S. Portugal et al., Host-mediated regulation of superinfection in malaria. Nat Med 17, 732–737 (2011).

10. Y. Sato, S. Ries, W. Stenzel, S. Fillatreau, K. Matuschewski, The Liver-Stage Plasmodium Infection Is a Critical Checkpoint for Development of Experimental Cerebral Malaria. Front Immunol 10, 2554 (2019).

11. H. Patel et al., Malaria blood stage infection suppresses liver stage infection via host-induced interferons but not hepcidin. Nat Commun 15, 2104 (2024).

12. A. F. Chora et al., Interplay between liver and blood stages of Plasmodium infection dictates malaria severity via gammadelta T cells and IL-17-promoted stress erythropoiesis. Immunity 56, 592–605 e598 (2023).

13. N. N. Ho, T. Tongogara, T. M. Tran, A. Kaushansky, E. K. Glennon, The case for exploring innate memory in Plasmodium infection. Curr Opin Microbiol 91, 102752 (2026).

14. A. Kaushansky, S. H. Kappe, Selection and refinement: the malaria parasite’s infection and exploitation of host hepatocytes. Curr Opin Microbiol 26, 71–78 (2015).

15. O. A. C. Lamers, B. M. D. Franke-Fayard, M. Roestenberg, J. M. M. Krol, The path from early- to late-liver stage arresting genetically attenuated parasites as a malaria vaccination strategy. NPJ Vaccines 10, 218 (2025).

16. M. G. Overstreet, I. A. Cockburn, Y. C. Chen, F. Zavala, Protective CD8 T cells against Plasmodium liver stages: immunobiology of an ’unnatural’ immune response. Immunol Rev 225, 272–283 (2008).

17. M. C. Aguirre-Botero, R. Amino, Tissue-dependent protection mechanisms of antibodies targeting Plasmodium sporozoites. Trends Parasitol 41, 853–867 (2025).

18. I. Tumwine-Downey et al., Antibody-dependent immune responses elicited by blood stage-malaria infection contribute to protective immunity to the pre-erythrocytic stages. Curr Res Immunol 4, 100054 (2023).

19. M. J. Stewart, R. J. Nawrot, S. Schulman, J. P. Vanderberg, Plasmodium berghei sporozoite invasion is blocked in vitro by sporozoite-immobilizing antibodies. Infect Immun 51, 859–864 (1986).

20. K. Vijayan et al., Antibody interference by a non-neutralizing antibody abrogates humoral protection against Plasmodium yoelii liver stage. Cell Rep 36, 109489 (2021).

21. F. Wunderlich, S. Al-Quraishy, M. A. Dkhil, Liver-inherent immune system: its role in blood-stage malaria. Front Microbiol 5, 559 (2014).

22. K. E. Lyke et al., Serum levels of the proinflammatory cytokines interleukin-1 beta (IL-1beta), IL-6, IL-8, IL-10, tumor necrosis factor alpha, and IL-12(p70) in Malian children with severe Plasmodium falciparum malaria and matched uncomplicated malaria or healthy controls. Infect Immun 72, 5630–5637 (2004).

23. G. L. Popa, M. I. Popa, Recent Advances in Understanding the Inflammatory Response in Malaria: A Review of the Dual Role of Cytokines. J Immunol Res 2021, 7785180 (2021).

24. I. C. Hirako et al., Uptake of Plasmodium chabaudi hemozoin drives Kupffer cell death and fuels superinfections. Sci Rep 12, 19805 (2022).

25. A. Kaushansky et al., Suppression of host p53 is critical for Plasmodium liver-stage infection. Cell Rep 3, 630–637 (2013).

26. T. M. Tran et al., A Molecular Signature in Blood Reveals a Role for p53 in Regulating Malaria-Induced Inflammation. Immunity 51, 750–765 e710 (2019).

27. P. Liehl et al., Host-cell sensors for Plasmodium activate innate immunity against liver-stage infection. Nat Med 20, 47–53 (2014).

28. P. Liehl et al., Innate immunity induced by Plasmodium liver infection inhibits malaria reinfections. Infect Immun 83, 1172–1180 (2015).

29. M. G. Netea, J. Quintin, J. W. van der Meer, Trained immunity: a memory for innate host defense. Cell Host Microbe 9, 355–361 (2011).

30. C. Marques-da-Silva et al., Type I interferons induce guanylate-binding proteins and lysosomal defense in hepatocytes to control malaria. Cell Host Microbe 33, 529–544 e529 (2025).

31. K. Tretina, E. S. Park, A. Maminska, J. D. MacMicking, Interferon-induced guanylate-binding proteins: Guardians of host defense in health and disease. J Exp Med 216, 482–500 (2019).

32. S. Christgen et al., The IFN-inducible GTPase IRGB10 regulates viral replication and inflammasome activation during influenza A virus infection in mice. Eur J Immunol 52, 285–296 (2022).

33. S. Martens et al., Disruption of Toxoplasma gondii parasitophorous vacuoles by the mouse p47-resistance GTPases. PLoS Pathog 1, e24 (2005).

34. J. G. Lu, P. Ji, S. W. French, The Major Histocompatibility Complex Class II-CD4 Immunologic Synapse in Alcoholic Hepatitis and Autoimmune Liver Pathology: The Role of Aberrant Major Histocompatibility Complex Class II in Hepatocytes. Am J Pathol 190, 25–32 (2020).

35. N. P. Riksen, M. G. Netea, Immunometabolic control of trained immunity. Mol Aspects Med 77, 100897 (2021).

36. M. A. Musrati et al., Infection history imprints prolonged changes to the epigenome, transcriptome and function of Kupffer cells. J Hepatol 81, 1023–1039 (2024).

37. A. Kang, M. D’Agostino, S. Afkhami, M. Jeyanathan, Z. Xing, Resident memory macrophages and trained innate immunity at barrier tissues. Elife 14, (2025).

38. Z. Wang et al., Tissue-resident trained immunity in hepatocytes protects against septic liver injury in zebrafish. Cell Rep 43, 114324 (2024).

39. J. L. Baratta et al., Cellular organization of normal mouse liver: a histological, quantitative immunocytochemical, and fine structural analysis. Histochem Cell Biol 131, 713–726 (2009).

40. A. Schepis et al., Elimination of intra-hepatocytic malaria parasites is driven by non-canonical autophagy but not nitric oxide production. iScience 28, 112052 (2025).

41. M. Charni-Natan, I. Goldstein, Protocol for Primary Mouse Hepatocyte Isolation. STAR Protoc 1, 100086 (2020).

42. W. Nahrendorf, A. Ivens, P. J. Spence, Inducible mechanisms of disease tolerance provide an alternative strategy of acquired immunity to malaria. Elife 10, (2021).

43. J. N. Crabtree et al., Lymphocyte crosstalk is required for monocyte-intrinsic trained immunity to Plasmodium falciparum. J Clin Invest 132, (2022).

44. C. L. Harding, N. F. Villarino, E. Valente, E. Schwarzer, N. W. Schmidt, Plasmodium Impairs Antibacterial Innate Immunity to Systemic Infections in Part Through Hemozoin-Bound Bioactive Molecules. Front Cell Infect Microbiol 10, 328 (2020).

45. K. Burleigh et al., Human DNA-PK activates a STING-independent DNA sensing pathway. Sci Immunol 5, (2020).

46. Y. Sohrabi et al., OxLDL-mediated immunologic memory in endothelial cells. J Mol Cell Cardiol 146, 121–132 (2020).

47. E. Kaufmann et al., BCG Educates Hematopoietic Stem Cells to Generate Protective Innate Immunity against Tuberculosis. Cell 172, 176–190 e119 (2018).

48. P. Heinke et al., Diploid hepatocytes drive physiological liver renewal in adult humans. Cell Syst 13, 499–507 e412 (2022).

49. R. Talwani, B. L. Gilliam, C. Howell, Infectious diseases and the liver. Clin Liver Dis 15, 111–130 (2011).

50. J. T. Robinson et al., Integrative genomics viewer. Nat Biotechnol 29, 24–26 (2011).

51. K. J. Livak, T. D. Schmittgen, Analysis of relative gene expression data using real-time quantitative PCR and the 2(-Delta Delta C(T)) Method. Methods 25, 402–408 (2001).

52. F. N. Watson et al., Cryopreserved Sporozoites with and without the Glycolipid Adjuvant 7DW8-5 Protect in Prime-and-Trap Malaria Vaccination. Am J Trop Med Hyg 106, 1227–1236 (2022).

53. Z. P. Billman, A. M. Seilie, S. C. Murphy, Purification of Plasmodium Sporozoites Enhances Parasite-Specific CD8+ T Cell Responses. Infect Immun 84, 2233–2242 (2016).

54. F. N. Watson et al., Ultra-low volume intradermal administration of radiation-attenuated sporozoites with the glycolipid adjuvant 7DW8-5 completely protects mice against malaria. Sci Rep 14, 2881 (2024).

55. B. K. Wilder et al., Anti-TRAP/SSP2 monoclonal antibodies can inhibit sporozoite infection and may enhance protection of anti-CSP monoclonal antibodies. NPJ Vaccines 7, 58 (2022).

56. D. Torre, A. Lachmann, A. Ma’ayan, BioJupies: Automated Generation of Interactive Notebooks for RNA-Seq Data Analysis in the Cloud. Cell Syst 7, 556–561 e553 (2018).

57. M. R. Corces et al., An improved ATAC-seq protocol reduces background and enables interrogation of frozen tissues. Nat Methods 14, 959–962 (2017).

58. J. D. Buenrostro, B. Wu, H. Y. Chang, W. J. Greenleaf, ATAC-seq: A Method for Assaying Chromatin Accessibility Genome-Wide. Curr Protoc Mol Biol 109, 21 29 21–21 29 29 (2015).

59. J. P. Smith, et al., PEPATAC: an optimized pipeline for ATAC-seq data analysis with serial alignments. NAR Genom Bioinform 3, lqab101 (2021).

60. M. Stolarczyk, V. P. Reuter, J. P. Smith, N. E. Magee, N. C. Sheffield, Refgenie: a reference genome resource manager. Gigascience 9, (2020).

61. H. M. Amemiya, A. Kundaje, A. P. Boyle, The ENCODE Blacklist: Identification of Problematic Regions of the Genome. Sci Rep 9, 9354 (2019).

62. S. Heinz et al., Simple combinations of lineage-determining transcription factors prime cis-regulatory elements required for macrophage and B cell identities. Mol Cell 38, 576–589 (2010).

63. Y. Chen, L. Chen, A. T. L. Lun, P. L. Baldoni, G. K. Smyth, edgeR v4: powerful differential analysis of sequencing data with expanded functionality and improved support for small counts and larger datasets. Nucleic Acids Res 53, (2025).

64. H. S. Kain et al., Liver stage malaria infection is controlled by host regulators of lipid peroxidation. Cell Death Differ 27, 44–54 (2020).

65. B. Tian et al., Interferon-Inducible GTPase 1 Impedes the Dimerization of Rabies Virus Phosphoprotein and Restricts Viral Replication. J Virol 94, (2020).

66. T. A. Kraus, L. Garza, C. M. Horvath, Enabled interferon signaling evasion in an immune-competent transgenic mouse model of parainfluenza virus 5 infection. Virology 371, 196–205 (2008).

